# METTL3 maintains epithelial homeostasis through m^6^A-dependent regulation of chromatin modifiers

**DOI:** 10.1101/2022.12.14.520484

**Authors:** Alexandra M. Maldonado López, Sijia Huang, Gina Pacella, Eun Kyung Ko, Hui Shen, Julian Stoute, Morgan Sinkfield, Amy Anderson, Stephen Prouty, Hua-Bing Li, John T. Seykora, Kathy Fange Liu, Brian C. Capell

## Abstract

The balance between epithelial stemness and differentiation requires the precise regulation of gene expression programs. Epitranscriptomic RNA modifications have been implicated in both epithelial development as well as cancers. However, the underlying mechanisms are poorly understood. Here, we show that deletion of the m^6^A methyltransferase, METTL3, impairs the m^6^A-mediated degradation of numerous mRNA transcripts encoding critical chromatin modifying enzymes, resulting in widespread gene expression abnormalities as well as both aberrant cutaneous and oral epithelial phenotypes *in vivo*. Collectively, these results offer new insights into a new layer of gene regulation within epithelial surface tissues and will inform future epitranscriptomic studies within epithelial cancer and developmental biology.

---

Stratifying epithelial tissues, such as the cutaneous and oral epithelia, provide an essential protective barrier throughout life which is constantly renewed by a stepwise differentiation program. In the skin, stem-like progenitor cells such as epidermal keratinocytes progress upwards to become terminally differentiated cells which make up a barrier against water loss and microbes (Baroni et al. 2012). This highly coordinated differentiation process requires precisely regulated spatiotemporal changes in gene expression (Miroshnikova et al. 2019), dysfunction of which can promote both neoplastic and inflammatory skin diseases (Rousselle 2016). One emerging area of gene regulation is that of epitranscriptomics (Novoa et al. 2017), or regulated RNA chemical modifications which offer an additional layer of gene regulation beyond modifications to DNA and histones (Lewis et al. 2017). While epitranscriptomic mechanisms have been implicated in a variety of processes ranging from stem cell differentiation to cancer, the role of the epitranscriptome in epithelial barrier tissues is poorly understood.

Amongst RNA modifications, *N*6-methyladenosine (m^6^A) is the most abundant chemical modification of messenger RNAs (mRNAs) and is enriched in exons, in the 3’ untranslated region (UTR) and near stop codons in human and mouse transcriptomes (Liu et al. 2020). m^6^A is catalyzed co-transcripabtionally on nascent pre-mRNA by an evolutionarily conserved, multicomponent writer complex with one known catalytic component, methyltransferase-like 3 (METTL3). METTL3-mediated m^6^A has been shown to regulate various aspects of mRNA metabolism, including both its stability and degradation by phase separation (Ries et al. 2019), as well as its translation (Lin et al. 2016; Slobodin et al. 2017; Zhou et al. 2018).

At the organismal level, METTL3-mediated m^6^A has been shown to regulate various aspects of cell development and homeostasis. For example, it facilitates rapid transcriptome turnover during cell differentiation to maintain homeostasis (Batista et al. 2014). Studies have shown that a reduction in m^6^A levels promotes cell differentiation, while an overall increase of methylation induces stemness and, therefore, suppresses cell differentiation (Zhao et al. 2017). Consistent with this, the loss of m^6^A via deletion of the writer catalytic subunit METTL3 promotes differentiation of mouse T cells (Li et al. 2017), and downregulation of METTL3 promotes upregulation of hematopoietic cell differentiation genes (Barbieri et al. 2017). In contrast, overexpression of METTL3 has been found in numerous cancers, including epithelial cancers such as oral and cutaneous squamous cell carcinomas (SCCs), and has been shown to be associated with stem-like properties and to accelerate the proliferation, invasion, and migration of SCC cells (Zhou et al. 2019; Zhao et al. 2020). Despite this, a deep mechanistic understanding of the functional roles of METTL3 and m^6^A is lacking. Therefore, in this study we combined both genome-wide methods and *in vivo* modeling to better understand the role of the METTL3-m^6^A epitranscriptome in epithelial tissues

## Results

### Loss of Mettl3 leads to grossly abnormal epidermal and tongue development

To better understand the role of the epitranscriptome in stratifying epithelia, we created mice with an epithelial-specific deletion of *Mettl3* by crossing mice floxed for *Mettl3* with mice carrying a keratin 14 (Krt14)- Cre (Krt14-Cre; *Mettl3*^fl/fl^ or “*Mettl3*-eKO”), as a whole-body deletion of *Mettl3* is embryonic lethal (Wu et al. 2018). *Mettl3*-eKO mice did not survive past one week of age, thus we harvested them for our experimental purposes on postnatal day 5 or 6 (P5 or P6). Phenotypically, *Mettl3*-eKO mice displayed pink skin at P5 and P6 that was strikingly different in comparison to wildtype (WT) controls (Fig. 1A), suggesting a potential failure to enter anagen, the period of active hair growth from postnatal days 1 to day 12 (Paus 1998; Fuchs et al. 2001; Plikus and Chuong 2008). Additionally, *Mettl3*-eKO mice were also about half the size of WT littermates (Fig. 1A). Notably, we observed that *Mettl3* heterozygotes weighed more than *Mettl3*-eKO mice, but less than WT control mice, suggesting that there is a correlation between weight loss with the loss of each allele of *Mettl3* (Fig. 1B). Together, these gross observations underscored how critical Mettl3 is for proper epithelial development.

**Figure 1.**
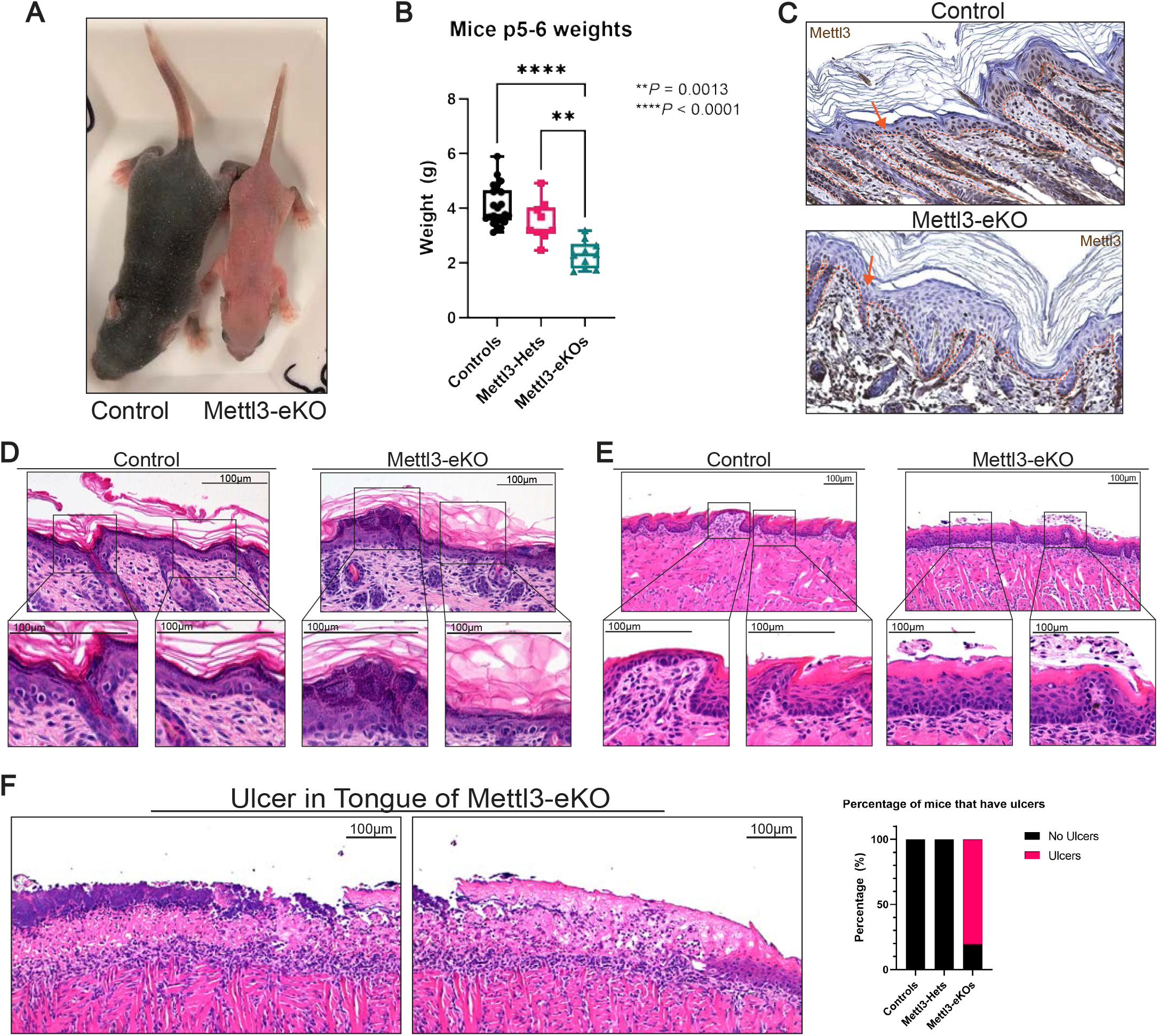
Loss of Mettl3 leads to grossly abnormal epidermal and tongue development. A) Postnatal day 6 (P6) Mettl3-eKO mice display a complete absence of hair and are half the size of WT control littermates. B) Mettl3-eKO mice weigh less than controls and Mettl3-heterozygotes (“Hets”). C) IHC of murine dorsal skin, the dotted orange line demarcates the the epidermis-dermis junction. The orange arrow signals the nucleus of the keratinocyte in which Mettl3 is found. Controls have Mettl3 present throughout the epidermis, but the staining is gone in the Mettl3-eKO epidermis. D) Mettl3-eKO skin displays malformed hair follicles, as well as loss of cell polarity in the basal epidermal cells along with thicker granular and cornified layers. E) Mettl3-eKO oral epithelium of the tongue has malformed filiform papillae and tastebuds. F) Mettl3-eKO tongues also display large ulcers in the posterior dorsal aspect.

We next examined the epithelial tissues histologically. Immunohistochemistry (IHC) confirmed a complete loss of Mettl3 throughout the epidermis (Fig. 1C). Using hematoxylin & eosin (H&E) staining, we saw that the skin of WT mice demonstrated normally formed epidermal layers with fully developed hair follicles and visible hair growth. In contrast, the *Mettl3*-eKO epidermis revealed several morphological abnormalities. For example, the interfollicular epidermis (IFE) displayed increased thickness of several epidermal layers, including stratum granulosum as well as the stratum corneum (Fig. 1D). In the basal layer of the epidermis, we found that the cell nuclei were not parallel to each other and displayed a loss of cell polarity, together suggesting potential defects in cell adhesion and proliferation (Watanabe et al. 2017). Beyond the IFE, and consistent with a previous study, we observed significant abnormalities in hair follicle morphogenesis (Fig. 1D) (Xi et al. 2020).

Since the Krt14-Cre system would also delete *Mettl3* in the oral epithelium (Su et al. 1993), we next examined the murine tongue. While WT control tongues demonstrated normal taste buds and filiform papillae across the dorsal lingual epithelium, *Mettl3*-eKO mice displayed severely impaired development of these features (Fig. 1E), consistent with previous work showing that loss of Mettl3 led to severe defects in taste bud development and abnormal epithelial thickening (Xiong et al. 2022). More surprisingly however, we observed that 80% of *Mettl3*-eKO mice were found to have full thickness ulcerations in the posterior aspect of the tongue, while none were present in WT controls, nor in *Mettl3*-heterozygotes (Krt14-Cre; *Mettl3*^+/fl^) (Fig. 1F), suggesting the *Mettl3*-eKO mice had impaired cell adhesion. Together, these results demonstrated that Mettl3 is a critical regulator of normal epithelial development and homeostasis.

### Mettl3 loss promotes significant transcriptional dysregulation

To begin to address the mechanisms behind how Mettl3 loss drove these dramatic phenotypic changes, we next performed RNA-seq from the dissected epidermis of *Mettl3*-eKO mice in comparison with their WT control littermates. Consistent with Mettl3’s established role in mRNA transcript regulation, as well as the numerous phenotypic abnormalities we were observing, *Mettl3*-eKO mice displayed profound transcriptional dysregulation, with almost 4,000 genes displaying significant differential expression upon *Mettl3* deletion (Fig. 2A)(Supplemental Table S1). In line with the increased epidermal thickening we observed in the Mettl3-eKO mice, Gene Ontology (GO) analysis of upregulated genes identified categories such as “Keratinization” and “Keratinocyte Differentiation”, along with specific transcripts like *Krt6a, Krt6b, Krt16*, and *Krt17* (Fig. 2B, C), all of which are associated with both thickened acral skin such as the human palm or sole, as well as in wounded and/or stressed keratinocytes after injury. Numerous other transcripts associated with keratinocyte terminal differentiation were significantly upregulated, including several small proline rich protein (*Sprr1a, Sprr2b, Sprr2d, Sprr2f, Sprr2g*) genes and several late cornified envelope (*Lce3a, Lce3b, Lce3d, Lce3f*) genes, as well as *Tgm1* and *Sfn*.

**Figure 2.**
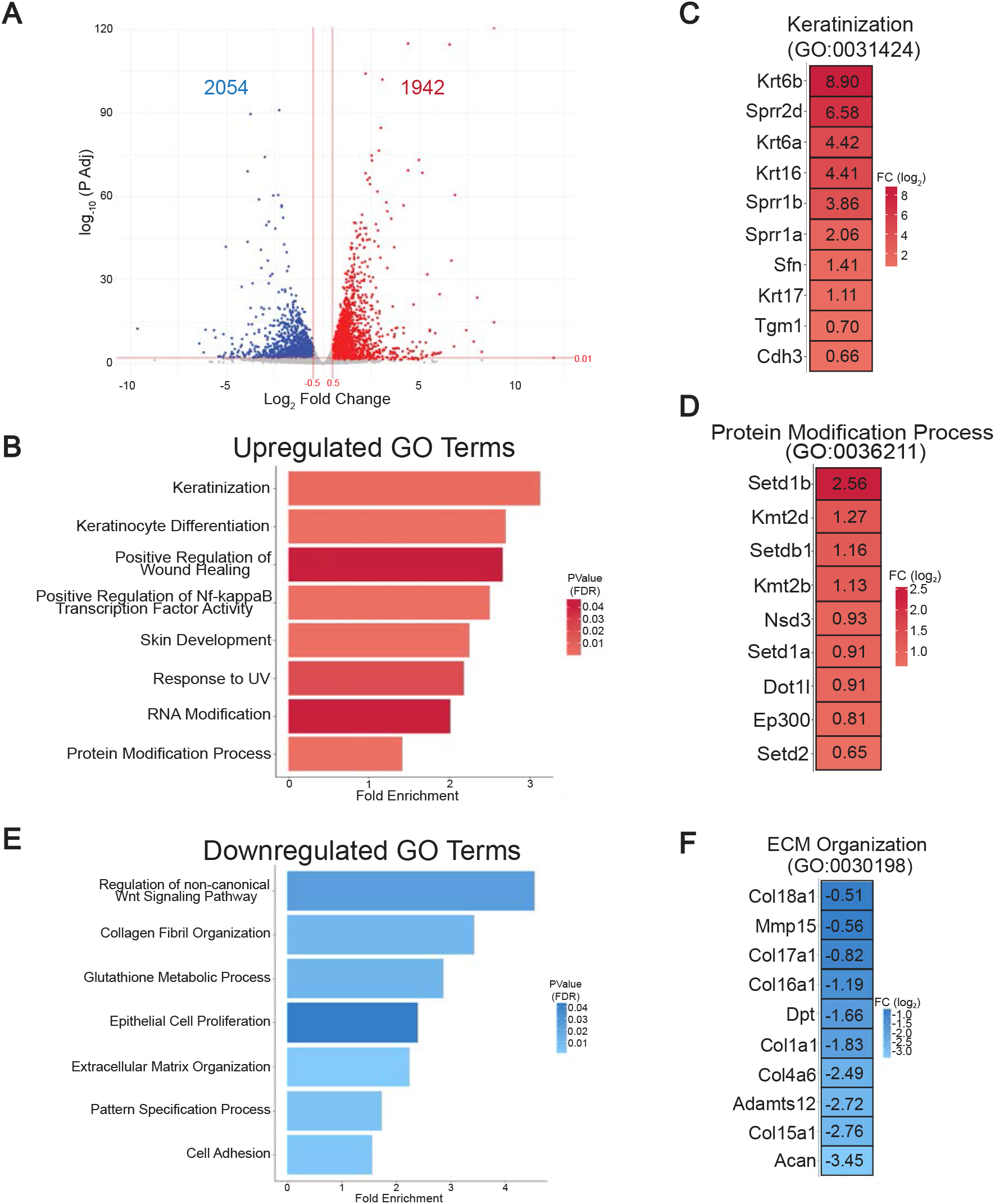
Mettl3 loss promotes significant transcriptional dysregulation. A) Bulk RNA-seq was done in Mettl3-eKO P6 mice (n = 3) vs Control littermates (n=3). A log2 Fold Change of ± 0.5 and an adjusted p-value of <0.01 was done to visualize the significant changes. B) Biological Process Terms Gene Ontology (GO:BP) Analysis of upregulated transcripts in Mettl3-eKO mice. C) Highly significant upregulated transcripts in Mettl3-eKO mice from the “Keratinization” category. D) Highly significant transcripts from the upregulated “Protein Modification Process” category. E) GO:BP Analysis of downregulated transcripts in Mettl3-eKO mice. F) Highly significant downregulated transcripts of the downregulated extracellular matrix (ECM) organization category.

Additionally, we noted enriched upregulated categories of “RNA Modification” and “Protein Modification Process”, which was particularly intriguing given the emerging role Mettl3 and m^6^A have been shown to play in the regulation of other epigenetic modifiers in a context-specific fashion (Kan et al. 2022). Transcripts identified by the “Protein Modification Process” were particularly notable for the broad representation of numerous chromatin modifiers, and in particular, histone methyltransferases. These included four H3K4 methyltransferases, Setd1a, Setd1b, Kmt2b (Mll2) and Kmt2d (Mll4), as well as methyltransferases for H3K36 (Setd2), H3K9 (Setdb1), H3K79 (Dot1l) (Fig. 2D). This was particularly interesting, as it has been shown previously that forced reduction of H3K4me3 by knockdown of KMT2B could reduce *TGM1* in normal human epidermal keratinocytes (NHEKs). In addition, knockdown of KMT2D caused a reduction of *SPRR2B*, together suggesting that these H3K4 methyltransferases may activate many genes involved in different stages of epidermal progenitor cell differentiation (Hopkin et al. 2012). In support of this, more recent work has demonstrated the importance of Kmt2d (Mll4) in promoting epidermal differentiation and tumor suppression *in vivo* (Egolf et al. 2021), while in contrast, enzymes that remove H3K4 methylation can suppress the expression of these terminal differentiation genes (Egolf et al. 2019).

Next, we performed GO analysis on the downregulated transcripts. In contrast to the enrichment of genes associated with differentiation seen amongst the upregulated transcripts, downregulated transcripts were enriched for pathways and categories typically associated with epithelial progenitor cells and more developmental pathways. These included categories such as “Regulation of non-canonical Wnt signaling”, “Epithelial Cell Proliferation”, “Extracellular Matrix (ECM) Organization”, “Collagen Fibril Organization”, “Cell Adhesion”, and “Pattern Specification Process” (Fig. 2E). All of these GO terms represent categories known to be important for the development and establishment of the basal layer of epithelial surface tissues like the skin epidermis and oral epithelium as well as hair follicle morphogenesis. For example, when examining the transcripts that made up these categories, we noted that numerous collagen genes were downregulated (Fig. 2F), including *Col17a1* (collagen XVII)(Fig. S1A, S1B), which facilitates epidermal-dermal attachment and regulates proliferation of epidermal stem cells in the interfollicular epidermis (IFE) (Natsuga et al. 2019). Intriguingly, mutations in human *COL17A1* are associated with junctional epidermolysis bullosa, a genetic blistering disease of the skin (Sproule et al. 2014; Hoffmann et al. 2019), while autoantibodies against collagen XVII are also known to be the primary cause of mucous membrane pemphigoid, a mucous membrane-dominated autoimmune subepithelial blistering disease that causes oral epithelial blistering (Xu et al. 2013). Collectively, these data present an overall picture whereby a loss of Mettl3 promotes a more differentiated, less progenitor cell-like phenotype. While the aberrant expression of numerous terminal differentiation transcripts may explain the thickened outer layers of the epidermis, the loss of transcripts involved in adhesion, including to the epithelial basement membrane, may offer insight into the loss of polarity observed in the epidermis, as well as the ulcerations observed in the tongue.

Finally, the broad upregulation of numerous chromatin modifying enzymes and histone methyltransferases demonstrated that a loss of Mettl3 may function to promote extensive gene expression alterations (and gross phenotypic abnormalities) through large-scale disruption of epigenetic and chromatin states.

### METTL3-mediated m^6^A dynamically regulates chromatin modifier mRNAs in epithelia

To gain further insight into the underlying mechanisms behind the phenotype in the *Mettl3*-eKO mice and how METTL3 and m^6^A function in epithelial tissues, we further examined human tissues such as the skin epidermis. Here were observed that METTL3 was present in the nuclei of keratinocytes, and its abundance decreased gradually from the basal stem-like progenitors to the terminally differentiated cornified layer (Fig. 3A). Consistent with this, we found that the global amount of m^6^A present in transcripts of differentiated keratinocytes was reduced in comparison to the progenitors upon *in vitro* differentiation of primary human keratinocytes (Fig. 3B). Together, these results suggested that higher levels of expression of METTL3 and m^6^A were associated with a more stem-like progenitor cell state, while reduced levels correlated with more differentiated epidermal cells and layers. This was consistent with our RNA-seq data which suggested that *in vivo* deletion of *Mettl3* promoted a more differentiated, less progenitor-like transcriptional phenotype.

**Figure 3.**
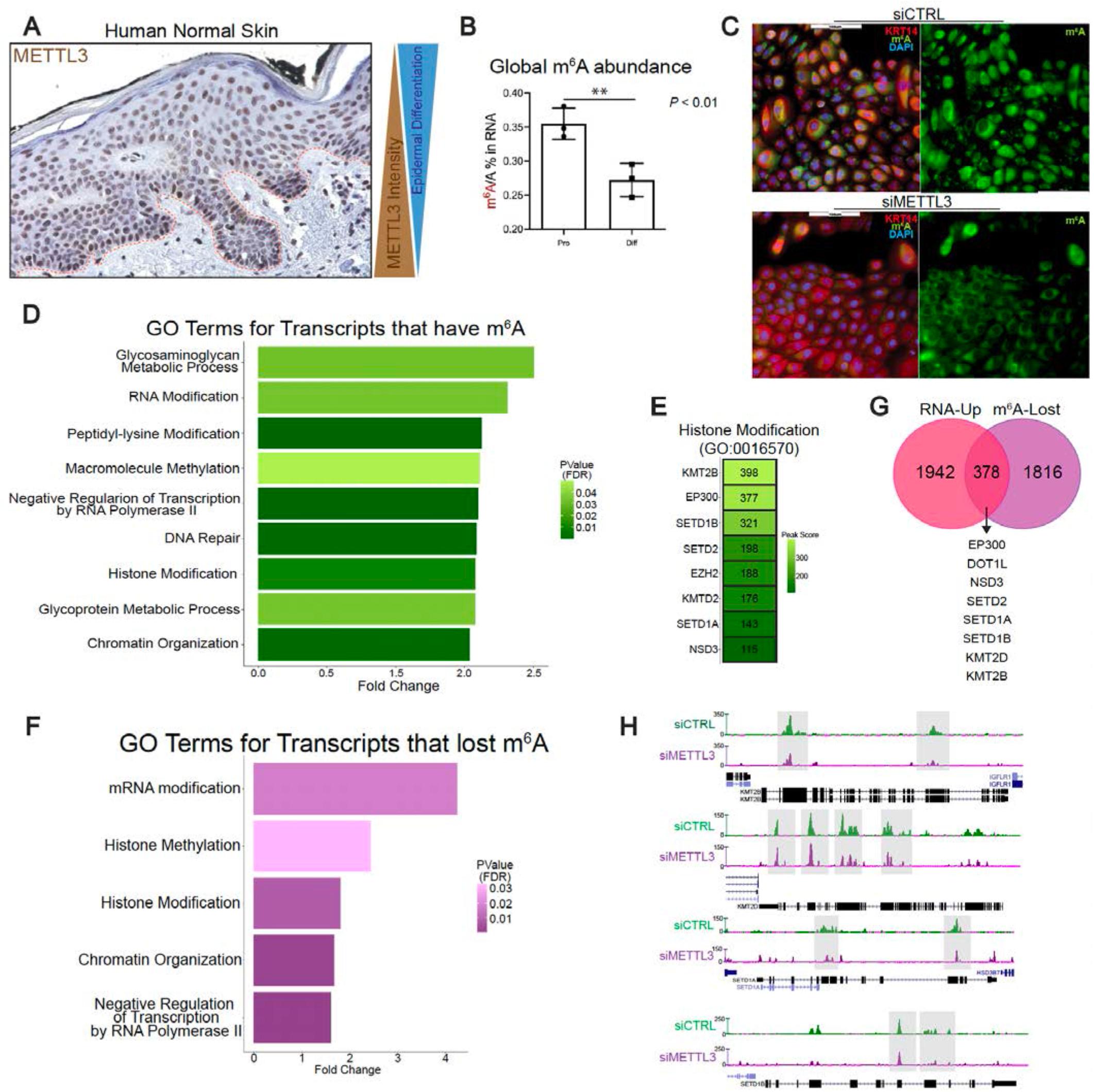
METTL3-mediated m^6^A dynamically regulates chromatin modifier mRNAs in epithelia. A) IHC of METTL3 in normal human skin, where above the dotted line is the epidermis. B) Quantitative analysis of the m^6^A level by LC-MS/MS done in proliferating vs differentiated neonatal human epidermal keratinocytes (NHEKs) demonstrates reduced global m^6^A with differentiation. C) METTL3 siRNA reduces visible m^6^A in NHEKs. D) GO analysis of transcripts which contain m^6^A peaks enriched in epidermal progenitors. E) Representative transcripts of the “Histone Modification” GO:BP category. F) GO analysis of transcripts which contain m^6^A peaks which are lost with siRNA depletion of METTL3. G) Overlap of the upregulated transcripts by RNA-seq (pink) and transcripts that lost m^6^A peaks with METTL3 depletion (purple). H) UCSC Genome Browser on Human (GRCh37/hg19) tracks from the m^6^A-seq analysis to visualize the decrease of m^6^A peaks in representative “Histone Modification” transcripts in control (green) and METTL3-depleted NHEKs (purple).

Next, utilizing this human epidermal progenitor cell model, we knocked down METTL3 via siRNAs and examined m^6^A (Fig. S2A). We verified a loss of m^6^A globally in these cells (Fig. 3C), and then performed m^6^A-sequencing (m^6^A-seq) to map m^6^A across all the mRNAs in the transcriptome. As previously reported, m6A was enriched in DRACH motifs (Fig. S2B). After calling common peaks across all human donor samples (Supplemental Table S2), we performed GO analysis to identify any enriched categories of m^6^A-modified mRNAs within the epidermal progenitors. Intriguingly, similar to the RNA-seq data in mice, our human model m^6^A-seq was enriched in categories representing epigenetic modifications, such as “Peptidyl-lysine Modification”, “Macromolecule methylation”, “Histone Modification”, and “Chromatin Organization” (Fig. 3D). Within these categories we found represented many of the same H3K4 methyltransferase transcripts including *SETD1A, SETD1B, KMT2B*, and *KMT2D* (Fig. 3E). To further validate these results, we examined if these methyltransferase transcripts lose m^6^A upon METTL3 depletion (Supplemental Table S2). To do this, GO analysis was performed on transcripts that displayed a significant reduction in m^6^A with siRNA METTL3 knockdown, and once again we saw that transcripts pertaining to the biological process categories of “Histone Methylation”, “Histone Modification”, and “Chromatin organization” were particularly enriched for a loss of m^6^A (Fig. 3F). Notably, all of these epigenetic modifiers were amongst the 378 transcripts that displayed both reduced m^6^A and increased mRNA abundance with METTL3 depletion (Fig. 3G). In contrast, only 152 transcripts that lost m^6^A also showed reduced mRNA expression (Fig. S2C). Direct visualization of these changes demonstrated that the intensity of m^6^A enrichment had decreased substantially in the transcripts of *SETD1A, SETD1B, KMT2B* and *KMT2D* in long internal exons, stop codons and/or 3’UTRs (Fig. 3H). These data suggested that METTL3-mediated m^6^A was directly regulating the abundance of these chromatin modifier mRNAs. In contrast, canonical keratinocyte genes associated with either the progenitor state (*COL17A1, COL7A1*) or the differentiated state (*LCE* genes, *SPRR* genes) that were dysregulated with *Mettl3* deletion did not display m^6^A enrichment or loss.

### Mettl3 loss increases mRNA half-life and enhances translation of chromatin modifiers

Given our data demonstrating that a loss of METTL3-mediated m^6^A on chromatin modifying enzyme transcripts was associated with their increased expression, we wanted to test whether METTL3 depletion was affecting the degradation of these transcripts. To do this, we employed a mRNA turnover assay (Ratnadiwakara and Anko 2018), which measures mRNA stability following the inhibition of transcription with Actinomycin D.

By inhibiting the synthesis of new mRNA, this allows for the measurement of mRNA decay by measuring mRNA abundance (Chen et al. 2008). Using this approach we measured the mRNA half-life of the *KMT2B, KMT2D, SETD1A* and *SETD1B* transcripts in epidermal progenitors following METTL3 depletion. Through this approach, we observed that *KMT2B, KMT2D* and *SETD1A* transcripts displayed evidence of a longer mRNA half-life upon METTL3 depletion (Figs. 4A and S3). This supported a model whereby the loss of METTL3-mediated m^6^A was leading to reduced degradation of chromatin modifier mRNA transcripts.

**Figure 4.**
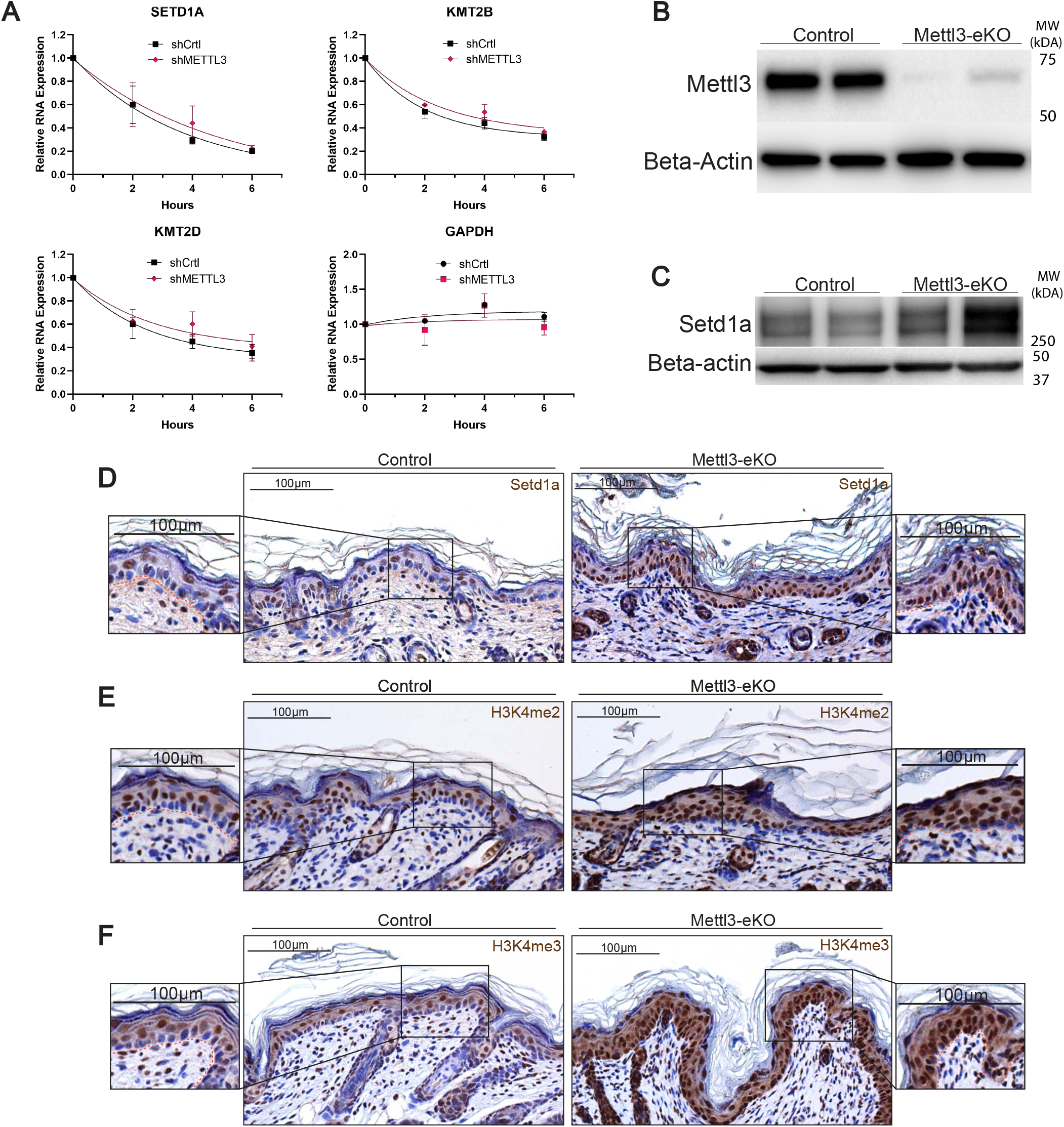
Mettl3 loss increases mRNA half-life and enhances translation of chromatin modifiers. A) mRNA turnover assay to measure the mRNA half-life transcripts in epidermal progenitors demonstrates prolonged half life of SETD1A, KMT2B and KMT2D transcripts with METTL3 depletion. GAPDH half-life is not altered by METTL3 knockdown. B) Immunoblotting of Mettl3 in control vs Mettl3-eKO P6 mouse epidermis. C) High molecular western blot for Setd1a in P6 mice epidermis. D) IHC for Setd1a in P6 mice epidermis. E) IHC for H3K4me2 in P6 mice epidermis. F) IHC for H3K4me3 in P6 mice epidermis.

Given these findings, we next asked whether these increases in mRNA transcript levels were also seen at the protein level, suggesting increased translation due to reduced degradation. We first examined SETD1A given its relatively abundant expression in human skin as per the Human Protein Atlas, as well as its essentiality for early mouse embryonic development (Bledau et al. 2014). Western blotting demonstrated an upregulation of Setd1a protein levels in *Mettl3*-eKO mice in comparison to WT controls (Fig 4B and 4C). Interestingly, upon staining the epidermis of *Mettl3*-eKO mice with Setd1a antibodies, we both confirmed the increased expression of the protein, but also noted that it was particularly enriched in the basal epidermal progenitors of the *Mettl3*-eKO mice in comparison to WT mice where it was largely absent from the basal cells (Fig 4D). As Setd1a is one of the major H3K4 histone methyltransferases in cells, we next interrogated levels of H3K4me2, which marks the gene body for activation, and H3K4me3, which primarily marks transcription start sites (Heintzman et al. 2009; Kranz and Anastassiadis 2020). Consistent with increased expression of multiple H3K4 methyltransferases including Setd1a, we found increased levels of both modifications throughout the epidermis. And similar to Setd1a expression, we also saw a particular enrichment of the modifications in the basal epidermal progenitors of *Mettl3*-eKO mice in comparison to WT controls (Fig. 4D and 4E).

Together, these data suggest a model whereby Mettl3-catalyzed m^6^A on H3K4 methyltransferase mRNAs promotes their degradation. In the absence of Mettl3, m^6^A is lost on these transcripts and their reduced degradation leads to their increased mRNA and protein abundance, ultimately resulting in both extensive epigenetic and phenotypic dysregulation (Fig. 5). Notably, enzymes that remove H3K4 methylation (i.e. the histone demethylase, LSD1) have been shown to repress differentiation (Egolf et al. 2019), while enzymes that catalyze H3K4 methylation (i.e. KMT2D/MLL4) have been shown to promote differentiation (Hopkin et al. 2012; Lin-Shiao et al. 2018; Egolf et al. 2021). Thus, this aberrant increased expression of these modifiers and the resulting increased H3K4 methylation in basal epidermal progenitors can promote the aberrant activation of more terminal differentiation genes and a loss of the progenitor state. This model is also consistent with emerging data demonstrating the existence of extensive crosstalk between epitranscriptomics and epigenetics (Kan et al. 2022).

**Figure 5.**
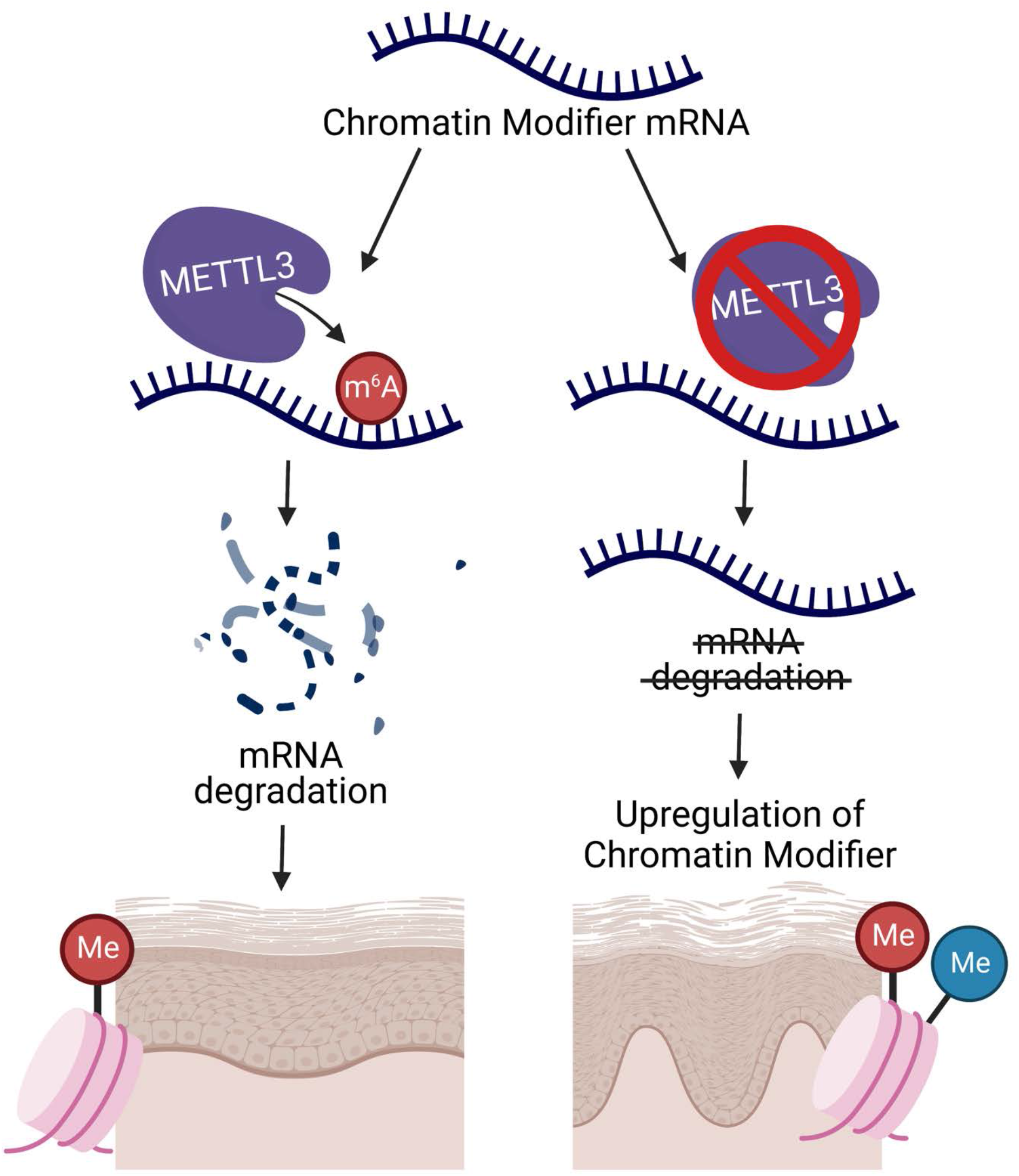
Schematic of METTL3-mediated m^6^A gene regulation in stratifying epithelial tissues. Loss of the METTL3-m^6^A epitranscriptome promotes the upregulation of chromatin modifiers by increasing their mRNA half-life and enhancing their transcripts’ translation. This upregulation of chromatin modifiers, and thus histone modifications, in turn, deregulate proper epidermal development and differentiation.

## Discussion

Our present study demonstrates that METTL3 is critical for the proper development of stratifying epithelial tissues and for maintaining the precise regulation of chromatin modifiers and histone modifications in these tissues. Building upon previous evidence of a role for Mettl3 in hair follicle development (Xi et al. 2020), our results demonstrate a novel role for Mettl3 and m^6^A in the direct regulation of the epigenome in stratifying epithelia. Specifically, we find that deletion of the m^6^A catalytic writer protein Mettl3 in the epithelial tissues of mice results in a striking phenotype in which the *Mettl3*-eKO mice are half the size and weight of their control littermates, are hairless, and unable to survive past a week while WT control littermates are healthy and thriving. In addition to the loss of normal hair follicle structures, basal layer keratinocytes in the IFE display a loss of polarity and adhesion along the basement membrane while the oral epithelium of the tongue displays striking ulcerations. In contrast, the more differentiated granular and the cornified layers are thicker. Consistent with these phenotypic alterations, RNA-seq demonstrated a broad loss of expression of genes involved in the epidermal progenitor state and adhesion to the basement membrane. These included adhesion genes like *Col7a1*and *Col17a1*, as well as genes associated with Wnt signaling and hair follicle stem cells (i.e. *Lef1, Wnt5a, Wnt7a, Krt15*). In contrast, upregulated genes included a variety of proteins associated with epithelial thickening and differentiation as well as numerous epigenetic regulators.

Mechanistically, we find that METTL3 depletion reduces m^6^A preferentially at chromatin modifying enzymes, and at H3K4 methyltransferases in particular. This results in specific significant increases in the mRNA levels of these transcripts due to impaired degradation (Fig. 5). While previous data has shown that increases in H3K4 methylation are associated with epithelial differentiation (Hopkin et al. 2012; Lin-Shiao et al. 2018; Egolf et al. 2021), our data demonstrates that METTL3 and m^6^A follow the opposite pattern and display reduced levels with differentiation. Our results further show that the observed changes in expression of differentiation genes (decreases in progenitor and epidermal basement membrane associated genes along with concomitant increases in terminal differentiation genes) are more likely secondary effects due to the widespread epigenetic dysregulation that occurs, in contrast to the direct m^6^A-mediated effects that loss of Mettl3 promotes on chromatin modifying transcripts.

Consistent with this model is a growing consensus in the field that a primary function of m^6^A is to promote mRNA degradation (Murakami and Jaffrey 2022). For example, recent evidence suggests that YTHDF epitranscriptomic reader proteins act redundantly to induce degradation of the same mRNAs (Zaccara and Jaffrey 2020). Additionally, these data are in line with extensive emerging evidence to show abundant interplay between epigenetics and the Mettl3-m^6^A epitranscriptome (Kan et al. 2022). For example, m^6^A has been shown to also promote the destabilization and degradation of chromatin modifying enzymes in neural stem cells (Wang et al. 2018). Furthermore, in addition to our results suggesting that H3K4 methylation was the primary modification to be regulated, other more recent studies have observed similar phenomena affecting heterochromatic marks such as H3K27me3 (Wu et al. 2020) and H3K9me2 (Li et al. 2020).

In summary, our results are the first discovery of how m^6^A regulates epigenetic regulators in epithelial tissues. They have revealed an entirely new layer of complexity to the epigenetic regulation of epithelial development and differentiation. While these results underscore the essential nature of normal Mettl3 and m^6^A function during epithelial development, important outstanding questions still remain, including what effects the post-development somatic loss of Mettl3 might have. Additionally, much remains to be discovered regarding the importance of epitranscriptomic regulation in epithelial diseases, particularly beyond cancers. However, even with the little we do know at this time, already potential therapeutic opportunities can be envisioned. For example, while METTL3 has been shown to be overexpressed in epithelial cancers like squamous cell carcinomas (Maldonado Lopez and Capell 2021), KMT2D/MLL4 is a known tumor suppressor that is frequently lost in those same cancers (Egolf et al. 2019). As there are numerous inhibitors of METTL3 being developed currently (Yankova et al. 2021; Berdasco and Esteller 2022), these data suggest that METTL3 inhibition could play a dually effective role of both inhibiting METTL3 and also increasing the expression of important tumor suppressors like KMT2D/MLL4. Given the great interest in and potential of drugs targeting epigenetic and epitranscriptomic regulators, we anticipate the results presented here will help inform those future studies in lieu of the epigenetic-epitranscriptomic crosstalk uncovered here.

## Materials and Methods

### Mice

All animal protocols were reviewed and approved by the Institutional Animal Care and Use Committee (IACUC) of the University of Pennsylvania. Mice were maintained on a C57BL/6 background. Mice carrying Mettl3 floxed alleles crossed with Krt14-Cre transgenic mice. Krt14-Cre; *Mettl3*^fl/fl^ (Mettl3-eKO) were considered mutants. Unless noted otherwise, littermates homozygous for the Mettl3 alleles of interest lacking Krt14-Cre were used as controls. The generation of *Mettl3*^fl/fl^ mice has been described elsewhere (Li et al. 2017).

### Genotyping

Polymerase chain reaction (PCR) was done using a Thermo Phire Animal Tissue PCR kit (F140WH). All experimental mice were an equal mix of males and females. The number of animals used per experiment is stated in the figure legends.

### Weight collection

Before euthanasia, mice at 6 days of age were measured for total body weight. Data figures are the composite of multiple litters of Mettl3-eKO or littermate controls. A one-way analysis of variance (ANOVA) was used to calculate the significance between groups when considered for genotype. All P are noted in figure legends and were considered significant if P < 0.05 and nonsignificant (NS) if P > 0.05.

### Mice RNA and protein extraction

Murine epidermis was dissociated from the dermis before the isolation of bulk RNA and protein. Following the euthanasia of 6-day-old mice, the skin was dissected, and the underlying fat pad was removed using a scalpel. The resulting tissue was floated dermis side down in dispase (5 U/ml; Corning) in PBS for 40 min at 37°C. The epidermis was then removed using a scalpel, and for RNA, the epidermis was flash-frozen in TRIzol and stored at −80°C until RNA isolation. RNA was extracted using a RNeasy kit (#74104, QIAGEN) at the same time and date for all mice belonging to a single experimental cohort (i.e., RNA-seq) regardless of the date of murine euthanasia to reduce batch effects and stored at −80°C. For protein, the epidermis was placed directly into cold PBS and centrifuged at 4°C for 5 min at 2500 rpm. To the resulting pellet, protein lysis buffer (Cell Signaling Technology) containing a protease inhibitor cocktail was added, and the mixture was homogenized, sonicated, rotated at 4°C for 10 min, and then centrifuged at 4°C at full speed for 10 min. Lysates were quantified using the Bradford assay. Frozen lysates were stored at −80°C.

### 2D Cell Culture

Proliferating primary Normal Human Epidermal Keratinocytes (NHEKs, or epidermal progenitors) cells were isolated from de-identified discarded neonatal foreskin provided by Core B - Skin Translational Research Core (STaR) of the Penn Skin Biology and Diseases Resource-based Center (NIH P30-AR069589). NHEK medium: filtered 50:50 mix of 1x keratinocyte serum-free medium (keratinocyte-SFM) supplemented with human recombinant epidermal growth factor and pituitary extract combined with medium 154 supplemented with human keratinocyte growth supplement (HKGS) and 1% 10,000 U/mL penicillin-streptomycin. Incubator: 37°C with 5% CO2. Proliferating NHEKs were cultured in NHEK medium under normal incubating conditions. Differentiating NHEKs arise from seeding proliferating NHEKs in NHEK medium for the first 24 hours after seeding and then changing the NHEK medium containing 1.22mM Calcium chloride (CaCl2) for 72 hours.

### siRNAs

Proliferating NHEKs cells cultured in penicillin-streptomycin free NHEK media and 24 hours later add the scrambled control siRNAs (siCTRL) or the siRNAs against METTL3 (siMETTL3) at 500nM dose for 72 hours under normal incubation conditions, changing to fresh penicillin-streptomycin free medium at 48 hours to prevent cell differentiation. Cells were harvested 72 hours after transfection.

### PCR

Complementary DNA was obtained using a high-capacity RNA-to-DNA kit (#4368814, Thermo Fisher Scientific). For quantitative real-time PCR, Power SYBR Green PCR Master Mix (#4367659, Thermo Fisher Scientific) was used. Quantitative real-time PCR data analysis was performed by first obtaining the normalized cycle threshold (CT) values (normalized to beta-actin and gapdh RNA), and the 2 − Ct method was applied to calculate the relative gene expression. ViiA 7 Real-Time PCR System was used to perform the reaction (Applied Biosystems). The average and SDs were assessed for significance using a Student’s t test. All P are noted in the figure legends and were considered significant if P < 0.05 and nonsignificant (NS) if P > 0.05.

### shRNAs

We generated stable fresh METTL3 knock down in proliferating NHEKs cell-lines and, after 2 or 3 passages to avoid compensatory mechanisms of loss of knockdown, we performed experiments.

### mRNA turnover assay

To assess the half-life of mRNA, Actinomycin D was used to treat the proliferating NHEK cells after shRNA knockdown. After the treatment, mRNA expression level at 0, 2, 4 and 6 h was analyzed via RT-qPCR following (Ratnadiwakara and Anko 2018).

### Quantitative analysis of the m^6^A level via LC-MS/MS

The PolyA+ RNA was extracted from total RNA using the Dynabeads mRNA Purification Kit (Ambion) following the manufacturer’s instructions. Purified RNA was digested and dephosphorylated to single nucleosides using Nucleoside Digestion Mix (NEB, M0649S) at 37 °C for 1 h. The detailed procedure was as previously described (Ontiveros et al. 2020). The nucleosides were quantified using retention time and the nucleoside-to-base ion mass transitions of 282.1 to 150.1 (m6A) and 268.0 to 136.0 (A). All quantifications were performed by converting the peak area from the LC-MS/MS to moles using the standard curve obtained from pure nucleoside standards running with the same group of samples. Then, the percentage ratio of m^6^A to A was used to compare the different modification levels.

### m^6^A-seq

Total RNA was extracted using the Qiagen RNeasy Mini kit (Cat No./ID: 74104). The following protocol was performed with 35-50ug of total RNA. mRNA was purified with Dynabeads mRNA purification kit (Invitrogen Catalog No. 61006). Fragmentation of the mRNA was performed using the Ambion RNA fragmentation reagents (Catalog #: AM8740). Then subjected to the RNA Clean and Concentrator-5 (Catalog No. R1013 from Zymo Research). mRNA quality was checked by Aligent BioAnalyzer 2100 using the RNA 6000 Pico Kit (Part number 5067-1513). One round of m^6^A immunoprecipitation was performed using the EpiMark N6-Methyladenosine Enrichment Kit (NEB #E1610S) and protocol. m^6^A-seq libraries were prepared using the NEBNext Ultra Directional RNA library preparation kit for Illumina (NEB #E7760). Library quality was checked by Agilent BioAnalyzer 2100 using the High Sensitivity DNA kit (Part number 5067-4626) and libraries were quantified using the Library Quant Kit for Illumina (NEB #7630). Libraries were then sequenced using a NextSeq500 platform (75-base-pair (bp) single-end reads). All kits were used following the manufacturer’s instructions.

### m^6^A-seq data processing

First, sequence quality was verified using fastqc. We used bowtie2 to map reads to the genome of mouse (mm10) with default parameters. Then, bedtools were used to remove the reads that contained low quality bases (MAPQ < 10) and undetermined bases. Mapped reads of IP and input libraries were processed by macs2 callpeak function, which identifies m^6^A peaks with bed or bam format that can be adapted for visualization on the UCSC genome browser. We called the peaks using the threshold of FDR < 0.05, no shifting model building, extension-size of 200bp and the removal of duplicates. HOMER is used for both the annotation of the called peaks and the de novo and known motif finding followed by localization of the motif with respect to peak summit. The differentially expressed peaks were selected with an absolute value of a fold change > 2 (and P < 0.05) by R package deseq2 (Love et al. 2014).

### RNA-seq

Total RNA was extracted using the Qiagen RNeasy Mini kit (Cat No./ID: 74104). The following protocol was performed with 1ug of total RNA. mRNA was purified using the NEBNext poly(A) mRNA magnetic isolation module. RNA-seq libraries were prepared using NEBNext Ultra Directional RNA library preparation kit for Illumina (NEB #E7760). Library quality was checked by Agilent BioAnalyzer 2100 using the DNA 1000 kit (Part number 5067-1504) and libraries were quantified using the Library Quant Kit for Illumina (NEB #7630). Libraries were then sequenced using a NextSeq500 platform (75-base-pair (bp) single-end reads).

### RNA-seq data processing

All RNA-seq was aligned using RNA STAR (Dobin et al. 2013) under default settings to Mus musculus GRCm38 FPKM (fragments per kilobase per million mapped fragments) generation and differential expression analysis were performed using DESeq2 (Love et al. 2014). Statistical significance was obtained using an adjusted p value (padj) generated by DESeq2 of less than 0.01.

### GO analyses

All GO analyses were performed using PANTHER at http://pantherdb.org/ to determine statistically overrepresented GO terms using Fisher’s exact test under the category “biological process.” P values for GO terms are false discovery rate statistics. The top eight plotted GO terms represent the GO terms with the highest fold enrichment under PANTHER’s default hierarchical clustering categorization. GO term figures are generated using ggplot2.

### Immunoblotting

Samples were separated by electrophoresis in 4 to 20% SDS–polyacrylamide gel electrophoresis gels with 30 ug per lane, transferred to a polyvinylidene difluoride membrane, and blotted with antibodies. Secondary horseradish peroxidase–conjugated secondary antibodies (Santa Cruz Biotechnology) and Amersham ECL Prime Western Blotting Detection Reagents (catalog no. RPN2232, GE Healthcare) were used for detection. For high molecular weight proteins, samples were separated by electrophoresis in 3 to 18% Tris-Acetate gel electrophoresis gels with 30 ug per lane, transferred to a polyvinylidene difluoride membrane, and blotted with antibodies. Secondary horseradish peroxidase–conjugated secondary antibodies (Santa Cruz Biotechnology) and Amersham ECL Prime Western Blotting Detection Reagents (catalog no. RPN2232, GE Healthcare) were used for detection.

### Histology

Mouse dorsal and ventral skin tissues were processed for histological examination by Core A - Cutaneous Phenomics and Transcriptomics (CPAT) Core of the Penn Skin Biology and Diseases Resource-based Center (NIH P30-AR069589) and mounted on frost-free slides. H&E staining was processed by Penn Skin Biology and Disease Resource-based Center Core A. A Leica DM6 B microscope was used to observe and capture representative images. Exposure times and microscope intensity were kept constant for all human samples and across mouse littermate comparisons.

### Immunohistochemistry

Tissue slides were baked for 1 hour at 65°C, deparaffinized in xylene, and rehydrated through a series of graded alcohols. After diH2O washes, slides were treated with antigen unmasking solution (1:100; SKU H-3300-250, Vector Laboratories) at 95°C for 12 min according to the manufacturer’s protocol. Hydrogen peroxide, blocking, primary antibody binding, HRP conjugated secondary antibody and 3, 3’- diaminobenzidine (DAB) were done using the Mouse and Rabbit Specific HRP/DAB IHC Detection Kit - Micro-polymer (ab236466, Abcam). Before the protein block, tissues were treated with 0.1% Triton X-100 diluted in 1X TBS for 5 minutes to unmask the antigens expressed in the nucleus. After overnight primary antibody incubation at 4°C and endogenous treatment with secondary anti-rabbit antibody at RT for 20 min, the staining was visualized with DAB with exposure times synchronized throughout all tissue samples within an antibody group for the exact same time. All slides were counterstained with hematoxylin (Hematoxylin QS, H-3404, Vector Laboratories) for 20s at RT, dehydrated in ethanol, cleared in xylene, and mounted with VectaMount (permanent mounting medium, H-5000, Vector Laboratories).

### Immunocytochemistry

Proliferating NHEKs with METTL3-KD were cultured in a chamber slide after being coated with 0.1% gelatin for 30 min at 37°C. After 48hrs, the chamber slides were fixed with 4% paraformaldehyde for 25 mins at RT and then washed with 1X DPBS. Chambers were permeabilized with 0.5% Triton X-100 in Dulbecco’s PBS for 10 min and blocked with 10% BSA in Dulbecco’s PBS for 2 hours in RT at gentle shaking. Primary antibodies diluted in 5% BSA in Dulbecco’s PBS were incubated O/N at 4°C with gentle shaking. Following fluorescent secondary antibody treatment 1 hr at RT in gentle shaking, the sections were mounted with ProLong Gold with 4′,6-diamidino-2-phenylindole (DAPI) (catalog no. P36935, Thermo Fisher Scientific).

### Statistical analyses

All statistical analyses were performed using R or GraphPad Prism 8. Details of each statistical test are included in Materials and Methods. Sample sizes and P values are included in the figure legends or main figures. Investigators were not blinded during experiments or outcome assessment.

## Supporting information

Supplemental Table S1 - RNA-seq

Supplemental Table S2 - m6A-seq

## Competing Interest Statement

The authors declare that they have no competing interests.

## Acknowledgements

Research reported in this publication was supported by the National Institute of Arthritis and Musculoskeletal and Skin Diseases (NIAMS) of the National Institutes of Health under Award Numbers (K08AR070289 and R01AR077615) to BCC, T32AR007465 to AMML. Additional support for this project was from R35GM133721 and R01HL160726 to KFL. The content is solely the responsibility of the authors and does not necessarily represent the official views of the National Institutes of Health.

Further support was provided by the Dermatology Foundation Stiefel Award for Skin Cancer to BCC and the Damon Runyon Cancer Research Foundation Clinical Investigator Award to BCC. This research was also supported by Cores A and B of the Penn Skin Biology and Diseases Resource-based Center, funded by NIAMS P30AR069589.

## Author contributions

Conceptualization: AMML, BCC

Data Curation: AMML, BCC

Formal Analysis: AMML, SH, JTS, KFL, BCC

Investigation: AMML, GP, EKK, HS, JS, MS, AA,

SP Supervision: BCC

Visualization: AMML,

SH Validation: SH, JTS, KFL, BCC

Writing – original draft: AMML, BCC

Writing – review & editing: AMML, SH, GP, EKK, HS, JS, MS, AA, SP, RAF, JTS, KFL, BCC

## Supplementary Figures and Methods are attached

## Notes

### Competing Interest Statement

The authors have declared no competing interest.

