## Supplemental Figures and Methods for "METTL3 maintains epithelial homeostasis through m^6^A-dependent regulation of chromatin modifiers"

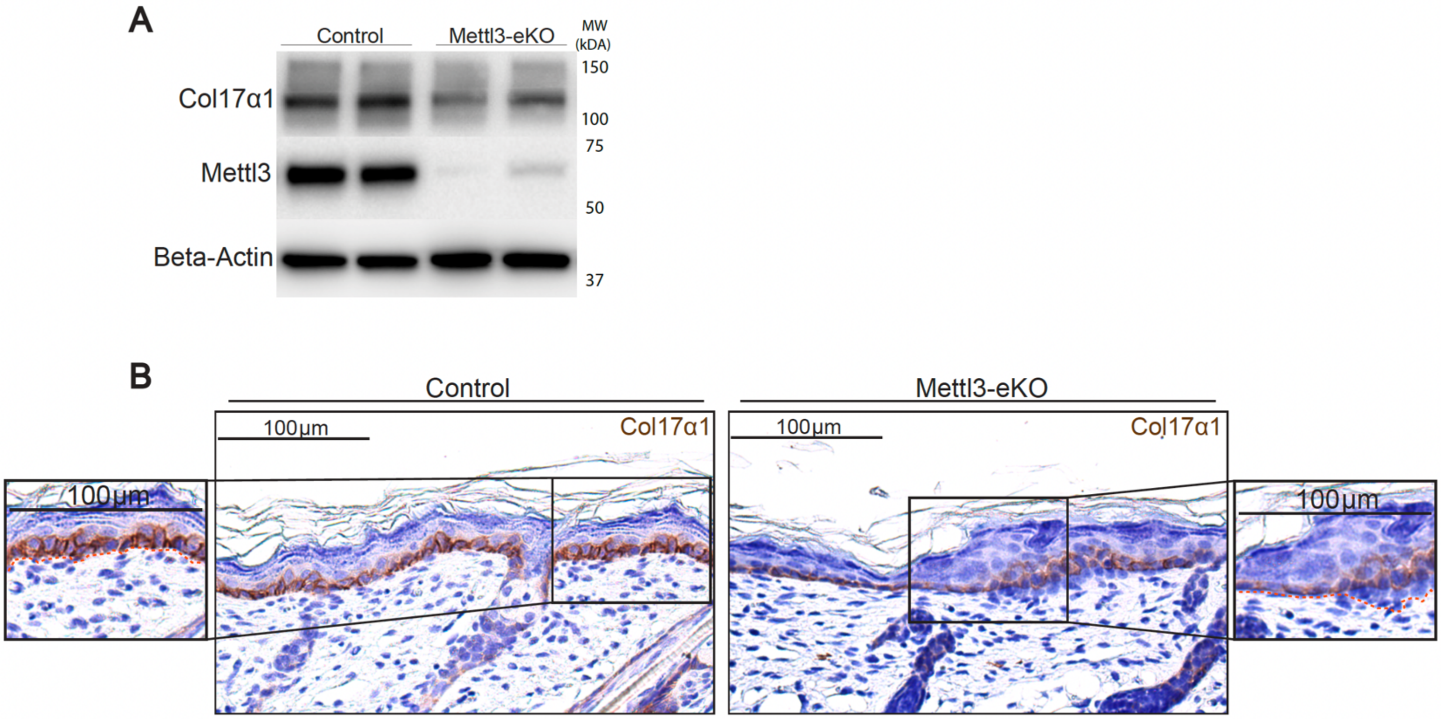


**Figure S1. Collagen 17 alpha 1 is decreased with deletion of *Mettl3*. (A)** Western blot for Col17a1 in P6 mice epidermis. **(B)** IHC for Col17a1 in P6 mice epidermis.

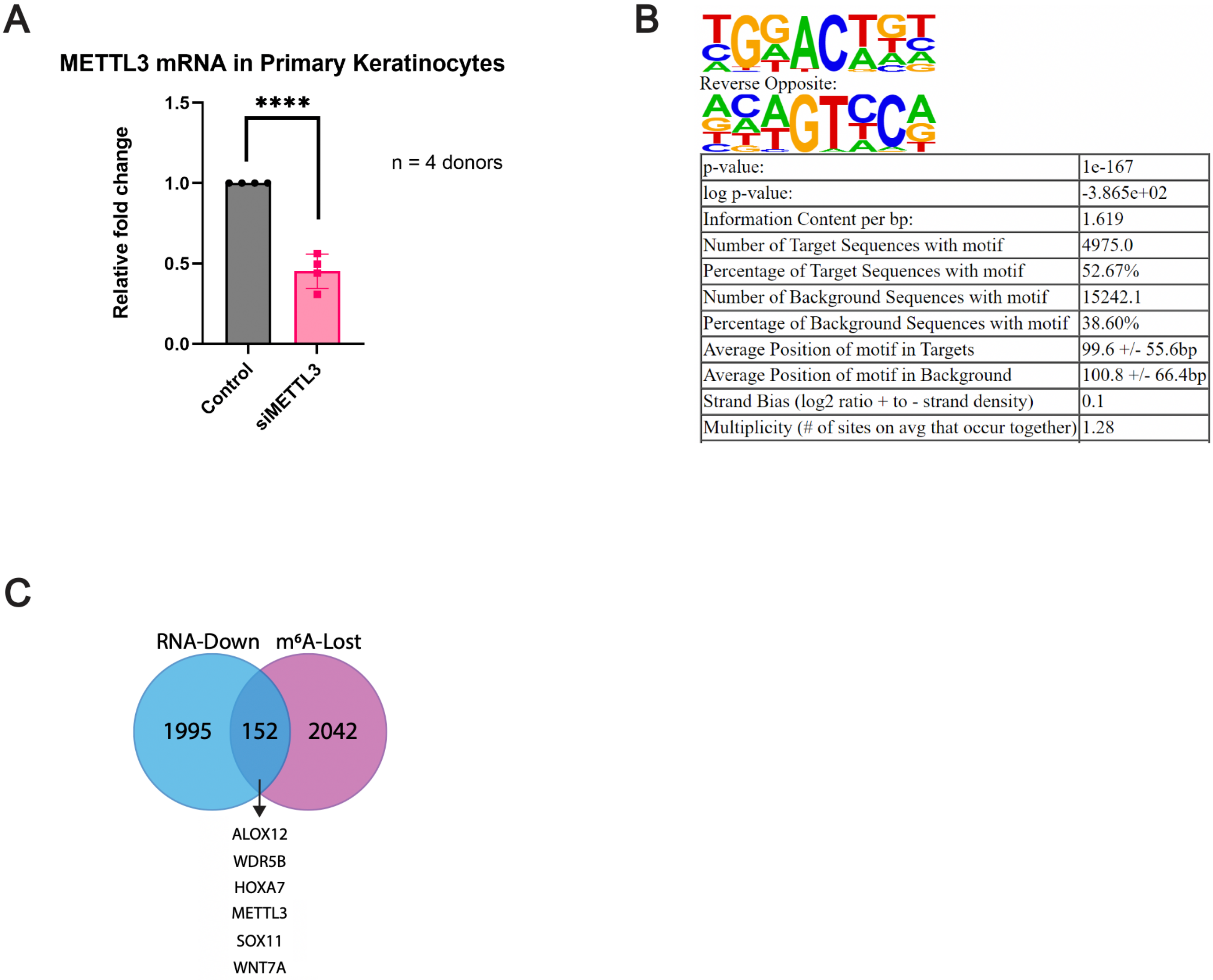


| | |
|---|---|
| p-value: | 1e-167 |
| log p-value: | -3.865e+02 |
| Information Content per bp: | 1.619 |
| Number of Target Sequences with motif | 4975.0 |
| Percentage of Target Sequences with motif | 52.67% |
| Number of Background Sequences with motif | 15242.1 |
| Percentage of Background Sequences with motif | 38.60% |
| Average Position of motif in Targets | 99.6 +/- 55.6bp |
| Average Position of motif in Background | 100.8 +/- 66.4bp |
| Strand Bias (log2 ratio + to - strand density) | 0.1 |
| Multiplicity (# of sites on avg that occur together) | 1.28 |

**Figure S2. METTL3-m$^6$A dynamics in epithelial tissues. (A)** RT-qPCR for METTL3 demonstrating knockdown with siMETTL3 treatment. **(B)** HOMER-identified enriched motifs for m$^6$A peaks in proliferating epidermal progenitors (normal human epidermal keratinocytes, or NHEKs). **(C)** Overlap of transcripts that demonstrate a concomitant loss of mRNA expression by RNA-seq (blue) as well as a significant reduction in m$^6$A enrichment by m$^6$A-seq (purple) with depletion of Mettl3.

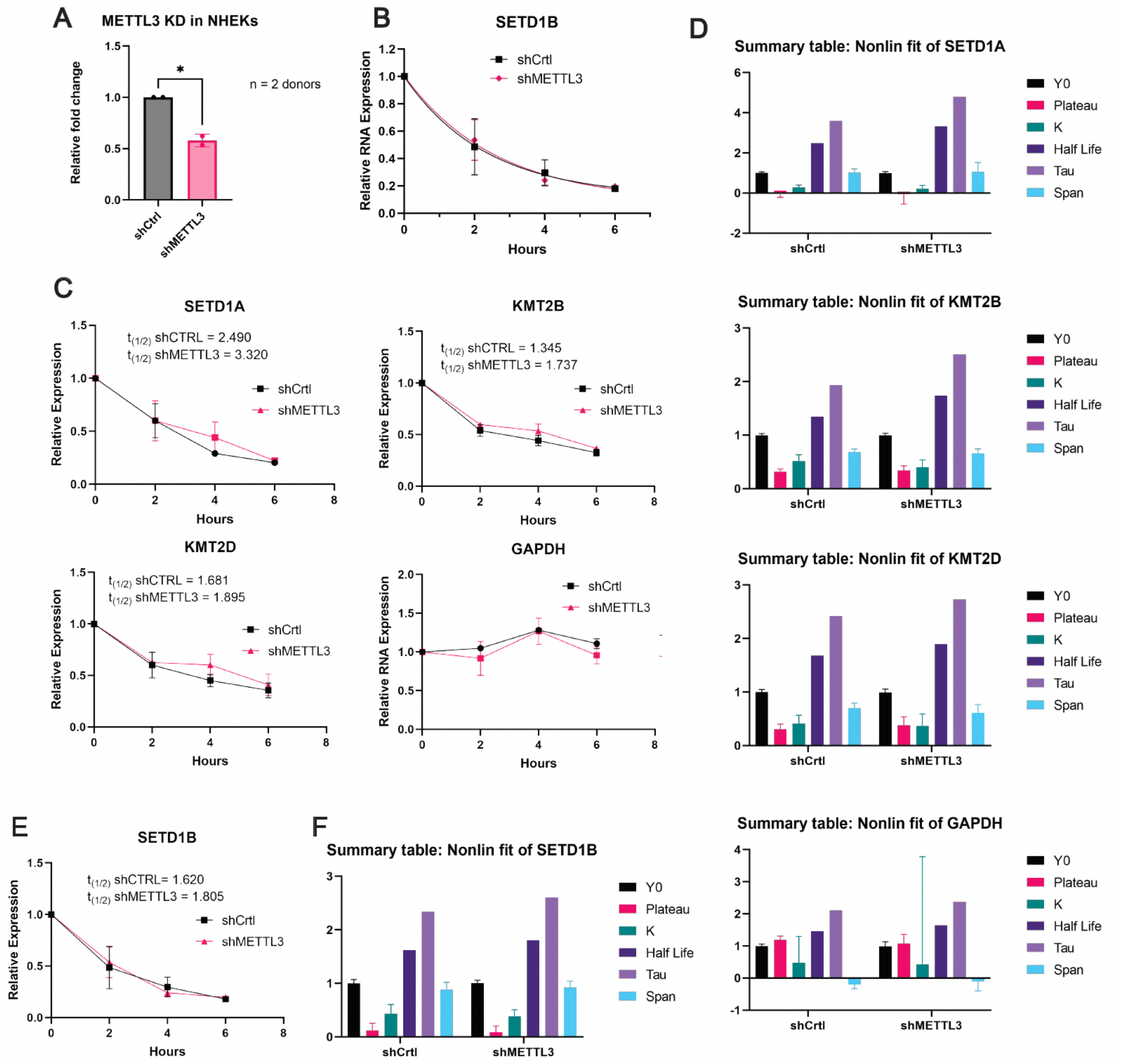


**Figure S3. mRNA half-life is increased with METTL3 loss. (A)** RT-qPCR for METTL3 shows knockdown with shRNA treatment. **(B)** mRNA turnover assay to measure the mRNA half-life of transcripts in proliferating epidermal keratinocytes (NHEKs) with and without METTL3 depletion. While the transcript half-life of *SETD1B* did not change appreciably when METTL3 is knocked down **(C)**, the calculated mRNA half-life (t1/2) of *SETD1A*, *KMT2B* and *KMT2D* transcripts increased when METTL3 was knocked down. *GAPDH*, a control housekeeping gene, was not altered by METTL3 knockdown. **(D)** Non-linear fit summary that was used to calculated mRNA half-life (t1/2) of *SETD1A*, *KMT2B*, *KMT2D* and *GAPDH*. **(E)** Calculated mRNA half-life (t1/2) of mRNA turnover assay of *SETD1B*. **(F)** Non-linear fit summary that was used to calculated mRNA half-life (t1/2) of *SETD1B*.

### Supplementary Materials

#### Antibodies

| Antibody | Company | Catalog Number | Application |
|---|---|---|---|
| Mettl3 | Abcam | ab195352 | Immunoblot, IHC |
| Setd1a | Invitrogen | PA5-78298 | Immunoblot, IHC |
| H3K4me2 | Abcam | ab7766 | IHC |
| H3K4me3 | Abcam | ab8580 | IHC |
| Cytokeratin 14 | Abcam | ab7800 | ICC |
| Beta-Actin | Cell Signaling | 4970S | Immunoblot |
| Collagen XVII | Novus Biologicals | NBP2-67316 | Immunoblot, IHC |
| $m^6A$ | Synaptic Systems | 202003 | ICC |
| Alexa Fluor 555 | Invitrogen | A31570 | ICC |
| Alexa Fluor 488 | Invitrogen | A21206 | ICC |
| Rb IgG HRP Linked | Cell Signaling | 7074S | Immunoblot |

#### siRNAs and shRNAs

| | |
|---|---|
| Accell Human *METTL3* siRNA - SMARTpool | Catalog ID: E-005170-00-0020 |
| Accell Non-targeting siRNA #1 | Catalog ID: D-001910-01-20 |
| Sigma METTL3 shRNA | SKU: TRCN0000034717 |
| Sigma pLKO.1-puro Non-Mammalian shRNA Control Plasmid DNA | SKU: SHC002 |

#### Primers (Genotyping and RT-PCR)

| | |
|---|---|
| *Mettl3* (mouse) | F' 5-AAGTGCTGCCATGTGAATGA-3<br>R' 5-TAAAGTGGAAAGGGTCAGTC-3 |
| *Keratin 14-Cre* (mouse) | F' 5-GAACCTGATGGACATGG-3<br>R' 5-AGTGCGTTCGAACGCTAGAGCCTGT-3 |
| *METTL3* (human) | F' 5-CAAGCTGCACTTCAGACGAA-3<br>R' 5-GCTTGGCGTGTGGTCTTT-3 |
| *SETD1A* (human) | F' 5-TTGCCATGTCAGGTCCAAAAA-3<br>R' 5-CGTACTTACGGCACATATCCTTC-3 |
| *SETD1B* (human) | F' 5-AGCGAGCTCCAGAACATGAC-3<br>R' 5-ATGGGTGTGAGGCATCTGTG-3 |
| *KMT2D* (human) | F' 5-ATCCTGGAGACACCCATCAG-3<br>R' 5-GACAGGCTCAGGGTCAGTG-3 |
| *KMT2B* (human) | F' 5-GGAGGAAGCAGCAAGCAGTA-3<br>R' 5-GCTCAGGTTTGGGGATTGT-3 |
| *Beta-Actin* (human) | F' 5-TGAAGTGTGACGTGGACATC-3<br>R' 5-GCAGGAGCAATGATCTTGAT-3 |

| | |
|---|---|
| *GAPDH* (human) | F' 5-ATCATCCCTGCCTCTACTGG-3<br>R' 5-GTCAGGTCCACCACTGACAC-3 |
